# Ventromedial Frontoinsular Connectivity is Associated with Long-term Smoking Behavior Change in Aging

**DOI:** 10.1101/2023.09.18.558264

**Authors:** Nagashree Thovinakere, Meishan Ai, Adrián E. Noriega de la Colina, Caitlin Walker, Giulia Baracchini, Jennifer Tremblay-Mercier, Sylvia Villeneuve, The PREVENT-AD Research Group, R. Nathan Spreng, Maiya R. Geddes

**Affiliations:** Department of Neurology and Neurosurgery, McGill University, Montreal, Canada; The Montreal Neurological Institute-Hospital, Montreal, Canada; Department of Psychology, Northeastern University, Boston, USA; Massachusetts Institute of Technology, Cambridge, USA

**Keywords:** Smoking, behavior change, ventromedial prefrontal cortex, anterior insula, resting-state fMRI, functional connectivity

## Abstract

A central question in the field of cognitive aging and behavioral neuroscience is what enables some individuals to successfully change their behavior more than others? Smoking is a significant risk factor for cognitive decline, particularly in vulnerable populations, including those who are at an elevated risk for Alzheimer’s disease (AD). Developing effective smoking reduction strategies is therefore a public health priority. The goal of the current study is to better understand the brain mechanisms underlying long-term smoking behavior change in cognitively normal, but at-risk, older adults. Neuroimaging and human lesional studies have implicated the insula and its functional network in subjective interoceptive awareness of cigarette craving and smoking-cue reactivity. We sought to characterize the extent to which anterior insular resting-state functional connectivity MRI predicted long-term smoking reduction (mean: 2.7 years, range 8 months – 4 years) using a seed-to-voxel approach. Twenty-three (18 women; 26% *APOE4* carriers; 61.5 years, SD = 3.7) cognitively unimpaired older individuals who smoked cigarettes at their baseline visit and have a first-degree family history of AD (at least one parent or multiple siblings affected) were included from a prospective longitudinal cohort, PREVENT-AD (Pre-symptomatic Evaluation of Experimental or Novel Treatments for Alzheimer Disease) in the current study. We found that reduced long-term smoking behavior was associated with diminished functional connectivity between bilateral anterior insula and ventromedial prefrontal cortex (vmPFC). In a second pre-registered replication study within a larger, independent sample of one hundred and eighteen cognitively normal older adults who smoked cigarettes at baseline from the UK Biobank (73 women; 27.9 % *APOE4* carriers; 60.3 years, SD = 2.7), we found that baseline diminished resting-state functional connectivity between anterior insula and vmPFC predicted long-term smoking reduction (mean 5.2 years; ranging from 3 years to 7 years). To our knowledge, this is the largest study to examine the neural substrates of long-term smoking cessation in human aging. Our results suggest that frontoinsular circuits may be a therapeutic target for smoking reduction and disease prevention in older adults at risk for AD.

## 1 Introduction

The global prevalence of Alzheimer’s disease (AD) and other dementias is projected to exceed 139 million by 2050 (Nichols et al., 2022), highlighting the urgent need to enhance disease prevention amidst an aging population. Targeting older individuals at risk for AD presents an opportunity for neuroprotection and disease risk reduction (McDade, 2022). In addition to age and *APOE ε4* carriership, modifiable vascular risk factors have emerged as significant contributors to cognitive decline and dementia (Gottesman & Seshadri, 2022; Livingston et al., 2020), with cigarette smoking accounting for the third highest population-attributable fraction of dementia cases (5.2%) after low educational attainment and hearing loss (Livingston et al., 2020).

Cigarette smoking is associated with diminished brain health in aging including increased white matter hyperintensities, acceleration of cerebral atrophy and poor cognitive functioning (e.g., memory and executive dysfunction) (Anstey et al., 2007; Gottesman & Seshadri, 2022; Livingston et al., 2020; M. C. Power et al., 2015; Rusanen et al., 2011). Smokers are at a higher risk of dementia compared to non-smokers, and dementia risk is reduced for individuals who quit or reduce smoking (Livingston et al., 2020). Quitting smoking, even later in life, has been linked to a reduced risk of cognitive decline in older adults (Choi et al., 2018). For example, an epidemiological study with over 50,000 individuals aged 60 and above found that quitting smoking for more than four years significantly lowered the risk of developing dementia in the subsequent eight years compared to those who continued smoking (Choi et al., 2018). Additionally, another study found that ex-smokers who maintained abstinence for at least three years had a similar risk of incident dementia as individuals who had never smoked (Lu et al., 2020).

There is evidence suggesting that reducing smoking, without necessarily quitting entirely, may also mitigate the risk of AD. Tyas et al. (2003) demonstrated a dose-dependent connection between smoking intensity and a heightened risk of AD and AD-related neuropathology, up to heavy smoking levels (>40.5 – 55.5 packs/year) (Tyas et al., 2003). Smoking reduction is a promising behavioural intervention strategy as a reduction in smoking may be a more attainable goal compared to complete cessation. Additionally, once achieved, it may further encourage attempts to quit without relapse. Currently however, the neurobehavioral mechanisms underlying smoking behavior change and the mechanisms of AD risk reduction itself in at-risk aging are completely unknown.

Despite the well-established benefits of smoking cessation, sustaining behavior change can be incredibly challenging. Long-term maintenance of behavior change, especially in the context of addictive behaviors, has proven to be a particular challenge due to the risk of relapse. Although some public health messaging and behavioral strategies have helped individuals quit smoking, and nicotine-dependent treatments are available, approximately 40 to 60 percent of individuals return to addictive substances within the first year after cessation (Etter & Stapleton, 2006; Janes et al., 2010; Stead et al., 2008). Understanding the mechanistic underpinnings of behaviour change maintenance has relied upon neuropsychological frameworks and cognitive-behavioural theories. Automatic processes, habits, self-efficacy expectations, and positive outcome expectations have been suggested to moderate smoking cessation maintenance (Baldwin et al., 2006; Dijkstra et al., 2007; Hertel et al., 2008). From a cognitive perspective, cognitive control processes, including response inhibition, working memory, and sustained attention have been associated with smoking behaviour change (Evans et al., 2018). Furthermore, to successfully maintain smoking cessation behaviour, executive functions, such as inhibitory control, goal-directed decision-making, and self-regulation are required (Hofmann et al., 2011; Kaplan & Berman, 2010). However, our understanding of the neural mechanisms underlying long-term maintenance of smoking reduction remains unknown, especially in an at-risk aging population. This investigation is especially relevant considering that the main barriers to successful smoking cessation and maintenance are withdrawal effects and cravings (Evans et al., 2018). A reduction in smoking could disrupt the neuroadaptive processes triggered by smoking cues, thereby reducing the likelihood of these cues inducing a desire to smoke in the future.

Human lesion and neuroimaging studies underscore the importance of the insula in smoking cessation and its role in conscious urges. These urges, which arise in response to specific cues, can be considered an emotional response involving a range of bodily responses, conscious feelings, and motivated behaviors triggered by objects of significant value to the individual (Naqvi & Bechara, 2010). Lesion studies have demonstrated a causal relationship between insular integrity, craving, and smoking cessation. For example, individuals with insular damage from a stroke showed abrupt decreased subjective cue-induced drug urges, less intense withdrawal symptoms, and less nicotine-seeking behavior compared to individuals with strokes in other brain regions (Naqvi et al., 2007). A study that applied lesion network mapping found that focal brain lesions resulting in addiction remission showed differential resting-state functional MRI (rs-fMRI) connectivity in a network involving the insula (Joutsa et al. 2022). These findings are consistent with task-based fMRI studies that have shown anterior insula activation in response to subjective cue-induced drug urges (Naqvi & Bechara, 2010). Heightened activation of the right anterior insula during a decision-making task, for instance, is associated with an increased likelihood of relapse to drug use (Claus et al., 2013; Janes et al., 2017; Luijten et al., 2011; Moeller et al., 2022; Namkung et al., 2017). Additionally, the anterior insula is crucial in predicting future reward and comparing present feelings with those from the past and the future, forming the basis of the involvement of this region in craving and drug dependence (Preuschoff et al., 2008). The anatomy and flexible functional connectivity profile of the anterior insula makes it a key hub within the salience network, allowing it to coordinate activity with the other major functional networks including the default mode network (DMN) and the central executive network (CEN) (Molnar-Szakacs & Uddin, 2022). Given its large scale connectivity, the anterior insula assumes a critical role in higher-order cognitive processes and goal-directed behaviours (Molnar-Szakacs & Uddin, 2022). Emerging evidence from resting-state network models propose that anterior insula is involved in directing attention towards either internal or external stimuli by mediating the dynamic activity between the other two large scale networks: DMN and the CEN (Sutherland et al., 2012).

In the present study, we aimed to assess whether altered baseline functional connectivity of the anterior insula predicted smoking behavior maintenance over the longer term in older adults at risk for AD. We applied rs-fMRI connectivity seed-to-voxel methods to address this guiding question in two separate studies. Rs-fMRI allows for the exploration of large-scale networks and their interactions, thus moving towards a systems-level understanding of brain function. First, we examined whether anterior insular connectivity predicted long-term smoking behavior change (mean = 2.7 years) in cognitively normal current smokers followed longitudinally in the PRE-symptomatic Evaluation of Experimental or Novel Treatments for Alzheimer’s Disease (PREVENT-AD) cohort at McGill University. We focused on this high-risk group as they most stand to benefit from smoking behavior change for disease prevention. We sought to ensure the robustness and generalizability of our results in an independent and larger sample. Based on the results of the first study, we preregistered the replication study design and hypothesis on the Open Science Framework platform (https://doi.org/10.17605/OSF.IO/BJV63). Next, we conducted the independent replication study utilizing the UK Biobank longitudinal cohort. In the replication study, our sample consisted of cognitively normal older adult (≥ 60 years) current smokers who were longitudinally followed for an average duration of 5.2 years. By employing this rigorous and comprehensive approach, we aimed to strengthen the validity of our conclusions and extend the significance of our findings to a broader aging population.

## 2 Methods

### 2.1 Participants in the PREVENT-AD Sample

Twenty-three (18 women) cognitively unimpaired older individuals who currently smoked cigarettes (mean age = 61.5 years, SD = 3.7) with a first-degree family history of AD (at least one parent or multiple siblings affected) were included from a prospective longitudinal cohort, PREVENT-AD. Out of 420 participants, our initial sample size consisted of 25 participants; two participants were removed from the final analyses for having poor-quality brain imaging data, resulting in a final sample size of 23 participants. Protocols and study procedures were approved by the McGill Institutional Review Board and/or Douglas Mental Health University Institute Research Ethics Board in accordance with the Declaration of Helsinki. Briefly, 399 cognitively healthy older individuals were enrolled between September 2011 and November 2017. Data from the present study are from Data Release 5.0. Inclusion criteria for the current cohort subsample were 1) age 60 years or older (or 55-59 years for individuals who were less than 15 years from the age of their relative at symptom onset); 2) no history of major neurological or psychiatric disease; 3) cognitively unimpaired at the time of enrollment; 4) reported being a current cigarette smoker at the baseline timepoint. Normal cognition was defined as a Clinical Dementia Rating (CDR) global of 0 and a Montreal Cognitive Assessment (MoCA) score of ≥26/30. Borderline individuals who did not fit the above criteria of normal cognition underwent a comprehensive diagnostic assessment by a trained neuropsychologist. All participants underwent longitudinal neuropsychological, AD genetic biomarker and serial MRI brain imaging assessments. All twenty-three participants who reported being current smokers at baseline, and for whom baseline structural and resting-state functional MRI (rs-fMRI) were available, were included in the present study. Brain imaging was obtained at baseline and self-reported smoking outcome data was obtained for two time-points: Baseline and follow-up (mean difference = 2.7 years, SD =1.56). Table 1 describes the demographic and neuropsychological information for the sample.

**Table 1.** Participant demographic characteristics. RBANS = Repeatable Battery for the Assessment of Neuropsychological Status. SD = Standard Deviation.

| **N=23** | **Mean±SD** |
| --- | --- |
| **Age at baseline (years)** | 61.5±3.7 |
| **Female sex (count and %)** | 18 (78.3%) |
| **Years of education** | 15.4±3.2 |
| **Baseline cigarettes/day** | 10.3±7.3 |
| **Follow-up cigarette/day** | 4.1±4.9 |
| **Number of quitters at follow-up (count and %)** | 8 (44.4%) |
| **APOE4 carrier status (positive count and %)** | 6 (26.1%) |
| **Age when started smoking (years)** | 16.2±2.6 |
| **Alcohol consumption (standard drinks/day)** | 1.1±0.8 |
| **Cognitive Function (RBANS Indices) at baseline** |  |
| **Immediate Memory Index** | 103.8±9.1 |
| **Visuospatial Index** | 100.1±13.7 |
| **Language Index** | 104.0±9.4 |
| **Attention Index** | 105.2±15.3 |
| **Delayed Memory Index** | 103.5±6.5 |
| **Total Score Index** | 103.9±8.9 |
| **Cognitive Function (RBANS Indices) at follow-up** |  |
| **Immediate Memory Index** | 108.7±12.5 |
| **Visuospatial Index** | 95.6±12.2 |
| **Language Index** | 98.3±9.1 |
| **Attention Index** | 107.4±13.3 |
| **Delayed Memory Index** | 104.1±8.5 |
| **Total Score Index** | 103.4±9.2 |

### 2.2 Participants in the Pre-registered UK Biobank Replication and Extension Study

One hundred and eighteen (118) cognitively unimpaired older individuals who currently smoked cigarettes (mean age = 60.32, SD = 2.70) were included from the UK Biobank cohort, a large population longitudinal cohort. This final sample size of 118 was a result after removing six participants for having poor brain-imaging data. Single nucleotide polymorphism (SNP) data for rs-429358 and rs-7412 were used to determine *APOE* genotypes. Inclusion criteria were 1) age 60 years or older; 2) no history of major neurological or psychiatric disease; 3) cognitively normal at the time of enrollment; 4) reported being current cigarette smokers at the baseline timepoint. Based on previous recommendations (Lyall et al., 2016). Normal cognition was defined as follows: performance scores on each of the cognitive test was converted into percentile rank, and the raw score corresponding to the 5^th^ percentile (or 95^th^, on tests where higher scores represented worse performance) was identified as the cut-off for impairment (Lyall et al., 2016). Imaging visit (Instance 2 of the UK-Biobank) was considered baseline timepoint, and the first repeat imaging visit (Instance 3 of the UK-Biobank) was considered the follow-up timepoint. Participants who reported being current smokers at baseline and for whom baseline structural and resting-state functional MRI (rsfMRI) data were available, were included in the present study. Brain imaging was obtained at baseline, and self-reported smoking outcome data was obtained for two timepoints: Baseline and follow-up (mean duration=5.2 years, SD=1.07). Table 2 describes the demographic and cognitive function information for the UK Biobank sample.

**Table 2.** Participant demographic characteristics for the UK Biobank replication cohort. SD = Standard Deviation.

|  |  |
| --- | --- |
| **N=118** | **Mean±SD** |
| **Age (years)** | 60.32±2.7 |
| **Female sex (count and %)** | 73 (61.9%) |
| **Years of education** | 14.6±3.2 |
| **Baseline cigarettes/day** | 11.3±5.9 |
| **Follow-up cigarette/day** | 6.1±3.9 |
| **Number of quitters at follow-up (count and %)** | 47 (39.8%) |
| ***APOE4* carrier status (positive count and %)** | 33 (27.9%) |
| **Age when started smoking (years)** | 15.2±1.6 |
| **Alcohol consumption (standard drinks/day)** | 1.8±0.6 |
| **Cognitive Function at baseline** |  |
| **Fluid Intelligence/Reasoning Score** | 6.9±1.8 |
| **Matrix Pattern Completion Score** | 7.8±2.1 |
| **Numeric Memory Score** | 6.9±1.3 |
| **Reaction Time (milliseconds)** | 597.3±96.1 |
| **Tower Rearranging Score** | 9.7±2.8 |
| **Trail Making Test, Part I (seconds)** | 212.9±47.8 |
| **Trail Making Test, Part II (seconds)** | 506.9±155.5 |
| **Cognitive Function at follow-up** |  |
| **Fluid Intelligence/Reasoning Score** | 6.8±2.0 |
| **Matrix Pattern Completion Score** | 7.7±2.0 |
| **Numeric Memory Score** | 6.9±1.2 |
| **Reaction Time (milliseconds)** | 601.9±95.3 |
| **Tower Rearranging Score** | 10.6±2.9 |
| **Trail Making Test, Part I (seconds)** | 233.6±59.6 |
| **Trail Making Test, Part II (seconds)** | 562.3±220.7 |

### 2.3 Smoking outcomes

#### 2.3.1 PREVENT-AD Sample

Smoking habits were assessed using the 7 items from the *Educoeur* questionnaire smoking sub-scale (number of cigarettes/day, age started smoking) (Goyer et al., 2013) at two timepoints, baseline and follow up (mean = 2.7 years, ranging from 8 months to 5 years). The behavioral outcome of interest (i.e., successful smoking behavior change) was defined as the difference between the number of cigarettes per day measured at follow-up compared to the number of cigarettes per day measured at baseline.

#### 2.3.2 UK Biobank Preregistered Replication Sample

Questions about smoking habits were taken from the touchscreen questionnaire on smoking habits (duration of smoking, number of cigarettes/day, age started smoking).

Individuals who report their current smoking status as “current smokers” at the first brain imaging timepoint were included in the study. The behavioral outcome of interest (i.e., successful smoking behavior change) is defined as the difference between the number of cigarettes per day measured at follow-up compared to the number of cigarettes per day measured at baseline (mean = 5.2 years, ranging from 3 years to 7 years).

### 2.4 Cognitive Indices

#### 2.4.1 PREVENT-AD Sample

Cognitive function was assessed using the Repeatable Battery for the Assessment of Neuropsychological Status (RBANS). This neuropsychological battery is a widely used and validated inventory. It includes five cognitive domains: Immediate Memory (i.e., list learning, story remembering), Visuospatial Ability (i.e., figure copy, line orientation), Language (i.e., picture naming, semantic fluency), Attention (i.e., digit span, coding) and Delayed Memory (i.e., list recognition, story recall and figure recall) (Randolph et al., 1998). To reduce test-retest effects, four different versions of the RBANS were used in follow up. The battery was administered annually to all participants enrolled in PREVENT-AD. Index scores are age-adjusted and have a mean of 100 with a standard deviation of 15.

#### 2.4.2 UK Biobank Preregistered Replication Sample

Brief cognitive tests were administered via touchscreen. The tests were designed specifically for UK Biobank but share some characteristics with other established tests of cognition. The following tests were included: Reaction Time, Numeric Memory, Fluid intelligence, Matrix pattern completion, Tower rearranging, and Trail making.

##### Reaction Time Test

Participants were instructed to quickly press a button with their dominant hand whenever a matching pair of symbols appeared on the screen. The test encompassed five practice trials followed by seven test trials. The metric analyzed was the average time (measured in milliseconds) taken to press the button. This value was derived from the four trials where a matching pair emerged (recorded in UK Biobank data field 20023). Higher scores indicated slower (i.e., worse) performance.

##### Numeric Memory Test

Participants were shown a sequence of numbers on the screen. After a brief pause, they were asked to recall and input the sequence in reverse order using a numeric keypad. Each number sequence was displayed for 2000 milliseconds, with an additional 500 milliseconds added for each digit in the sequence. A delay of 3000 ms occurred between clearing the screen and activating the response keypad. The test began with a sequence length of two digits and progressed to longer sequences, increasing by one digit each time, up to a maximum length of 12 digits. The test ended either after five consecutive incorrect responses for two-digit sequences or after two consecutive incorrect responses for sequences of three digits or more. The analysis score was based on the longest correctly recalled sequence length (recorded in UK Biobank data field 4282), with higher scores indicating better performance.

##### Fluid Intelligence

Thirteen questions were presented in succession on a touchscreen for the Fluid Intelligence test. Participants could respond at their own pace within a two-minute time frame. Responses were chosen from a set of multiple-choice options. Any questions not attempted within the two-minute period received a score of zero. The analysis score represented the total number of correct answers, ranging from 0 to 13 (UK Biobank data field 20016), with higher scores indicating better performance.

##### Matrix pattern completion

Non-verbal fluid reasoning was assessed through an adapted version of the COGNITO Matrices test (De Roquefeuil Guilhem, 2014). Participants were presented with matrix pattern blocks, each with a missing element. Their task was to select the element that completed the pattern best from a range of choices. The items varied in difficulty across a total of 15 items, and the analysis score was the count of correctly answered items within a 3-minute interval.

##### Tower rearranging

To gauge planning abilities, an adapted form of the One-touch Tower of London test (Hampshire et al., 2013) was employed. Participants were shown illustrations featuring three differently colored hoops positioned on three pegs. Their task was to determine the number of moves needed to rearrange the hoops to a specified configuration. The analysis score was the tally of correctly answered items within a 3-minute period.

##### Trail Making Test

The UKB Trail Making Test (TMT) is a digital adaptation of the widely recognized Halstead-Reitan Trail Making Test (Reitan & Wolfson, 1985), often regarded as an assessment of executive function. The test comprised two parts. Part A displayed numbers 1-25 arranged pseudo-randomly on the screen, requiring participants to touch them in numerical order. Part B presented numbers 1-13 and letters A-L arranged pseudo-randomly, necessitating participants to alternate between touching numbers in numeric order and letters in alphabetical order (e.g., 1-A-2-B-3-C). Participants were instructed to complete each part as swiftly and accurately as possible. Incorrect responses prompted a red screen flash, requiring participants to correct their answer before proceeding. The analysis score represented the time taken, in seconds, to complete each part.

### 2.5 MRI data acquisition

#### 2.5.1 PREVENT-AD Sample

All participants underwent an MRI scanning session in a 3T Magnetom Tim Trio (Siemens) scanner using a Siemens standard 12-channel head coil. T1-weighted structural images were obtained using a GRE sequence with the following parameters: TR = 2300 ms; TE = 2.98 ms; flip angle = 9°; FOV = 256 × 240 × 176 mm; voxel size = 1 × 1 × 1 mm. For resting state functional MRI scans, two consecutive functional T2*-weighted runs were collected with eyes-closed using a blood oxygenation level-dependent (BOLD) sensitive, single-shot echo planar sequence with the following parameters: TR = 2000 ms; TE = 30 ms; flip angle=90°; FOV= 256 × 256 mm; voxel size=4×4×4 mm; 32 slices; 150 volumes, acquisition time = 5min4s per run.

#### 2.5.2 UK Biobank Preregistered Replication Sample

Details of the image acquisition and processing are available on the UK Biobank Protocol (http://biobank.ctsu.ox.ac.uk/crystal/refer.cgi?id=2367), and Brain Imaging Documentation (http://biobank.ctsu.ox.ac.uk/crystal/refer.cgi?id=1977). Briefly, all brain MRI data were acquired on a single standard Siemens Skyra 3T scanner with a standard Siemens 32-channel RF receiver head coil, with the following parameters: TR = 2000 ms; TI = 800 ms; R =2; FOV = 208 × 256 × 256 mm; voxel size = 1 × 1 × 1 mm.For resting state functional MRI scans, two consecutive functional T2*-weighted runs were collected with eyes-closed using a blood oxygenation level-dependent (BOLD) sensitive, single-shot echo planar sequence with the following parameters: TR = 735 ms; TE = 39 ms; flip angle=52°; FOV 88 x 88 x 64 matrix; resolution=2.4×2.4×2.4 mm; 490 volumes, acquisition time = 6min per run.

### 2.6 Resting-state functional MRI Data Preprocessing

Preprocessing of raw resting state functional images from both cohorts was completed using an identical approach. This included application of fMRIprep (version 20.2.4) (Esteban et al., 2019). For each of the BOLD runs per subject, the following preprocessing was performed: First, the T1w reference is skull-stripped using a Nipype implementation of the antsBrainExtraction.sh tool (ANTs). A B0-nonuniformity map (or fieldmap) was estimated based on a phase-difference map calculated with a dual-echo Gradient-Recall Echo (GRE) sequence, which was then co-registered to the target echo-planar imaging (EPI) reference run and converted to a displacements field map. A distortion-corrected BOLD EPI reference image was constructed and registered to the T1-weighted reference using a boundary-based approach (*bbregister, Freesurfer*). Rigid-body head-motion parameters with respect to the BOLD EPI reference were estimated (*mcflirt, FSL 5.0.9*) (Jenkinson et al., 2002) before any spatiotemporal filtering was performed. BOLD runs belonging to the single band acquisition sessions were slice-time corrected (using *3dTshift, AFNI 20160207*). The BOLD time-series were resampled onto the fsaverage surface (FreeSurfer reconstruction nomenclature). The BOLD time series (including slice-timing correction when applied) were resampled into their original, native space by applying a single, composite transform to correct for scan-to-scan head motion and susceptibility distortions. The BOLD time series were resampled into the standard space using the MNI152 template. Several confounding time-series were calculated based on the preprocessed BOLD: framewise displacement (FD), DVARS and three region-wise global signals. FD and DVARS are calculated for each functional run, both using their implementations in Nipype (following the definitions by Power et al., 2014). The three global signals were extracted within the cerebrospinal fluid (CSF), the white matter (WM), and the whole-brain masks. Following previous recommendations to achieve stable results, functional scans were spatially smoothed using a 6 mm full width at half maximum (FWHM) Gaussian smoothing kernel. During the outlier detection step, acquisitions with framewise displacement above 0.9 mm or global BOLD signal changes above 5 standard deviations were flagged as outliers using the Artefact Detection Tool (www.nitrc.org/projects/artifact_detect).

Additionally, scan-to-scan mean head motion (framewise displacement) was used as a covariate of non-interest in all second level analyses (mean head motion= 0.2mm, SD=0.1mm). Head motion is a known important potential confound as it produces systematic and spurious patterns in connectivity, and is accentuated in AD and cognitively typical aging populations (Power et al., 2012). Critically, we did not identify a relationship between the mean head motion parameter and either smoking behavioral change or individual functional connectivity (all *p* > 0.05). Two participants were removed from the PREVENT-AD sample final analysis for having > 30 scan volumes flagged, leading to the final sample size of 23 participants. Similarly, six participants were removed from the UK-Biobank cohort for the same reason, resulting in the final sample size of 118 participants. This cut off was determined based on preserving at least 5 min of scanning time (Van Dijk et al., 2010). A set of physiological regressors were extracted to allow for principal component-based noise correction (CompCor41). Principal components were estimated for the temporal and anatomical CompCor variants. The following nuisance parameters were regressed out from the functional time series using ordinary least-squares regression: six motion parameters and their derivatives, global signal, framewise displacement (Power et al., 2014).The first six noise components were estimated by aCompCor (Behzadi et al., 2007) and included polynomial trends up to second order. Temporal band-pass filtering (0.008– 0.09 Hz) was applied using the CONN toolbox to remove physiological, subject-motion and outlier-related artefacts (version 20.c, https://www.nitrc.org/projects/conn/) (Whitfield-Gabrieli & Nieto-Castanon, 2012).

### 2.7 Seed-to-voxel Functional Imaging Data Analysis

To investigate the relationship between brain connectivity and long-term smoking change behavior outcome, seed-to-voxel analyses were implemented identically in the PREVENT-AD sample and the UK Biobank as a replication and extension sample. Resting-state functional brain connectivity analyses were carried out using the CONN toolbox (version 20.c, https://www.nitrc.org/projects/conn/) (Whitfield-Gabrieli & Nieto-Castañón, 2012). The average time series in the anatomically defined region of interest (ROI), bilateral anterior insula (aI) was extracted (Figure 1a). The ROI was defined based on the Harvard-Oxford atlas. Product-moment correlation coefficients were computed between the average time series within the ROI and the time series within all other voxels in the whole brain and converted to normally distributed Fisher transformed z-scores prior to entering the second-level general linear model. Individual change in smoking was entered as a covariate of interest in the second-level analysis, controlling for nuisance variables including age, sex, scan-to-scan mean head motion, *APOE4* carrier status and baseline smoking amount (number of cigarettes/day) in a general linear model for the bilateral aI.

**Figure 1.**
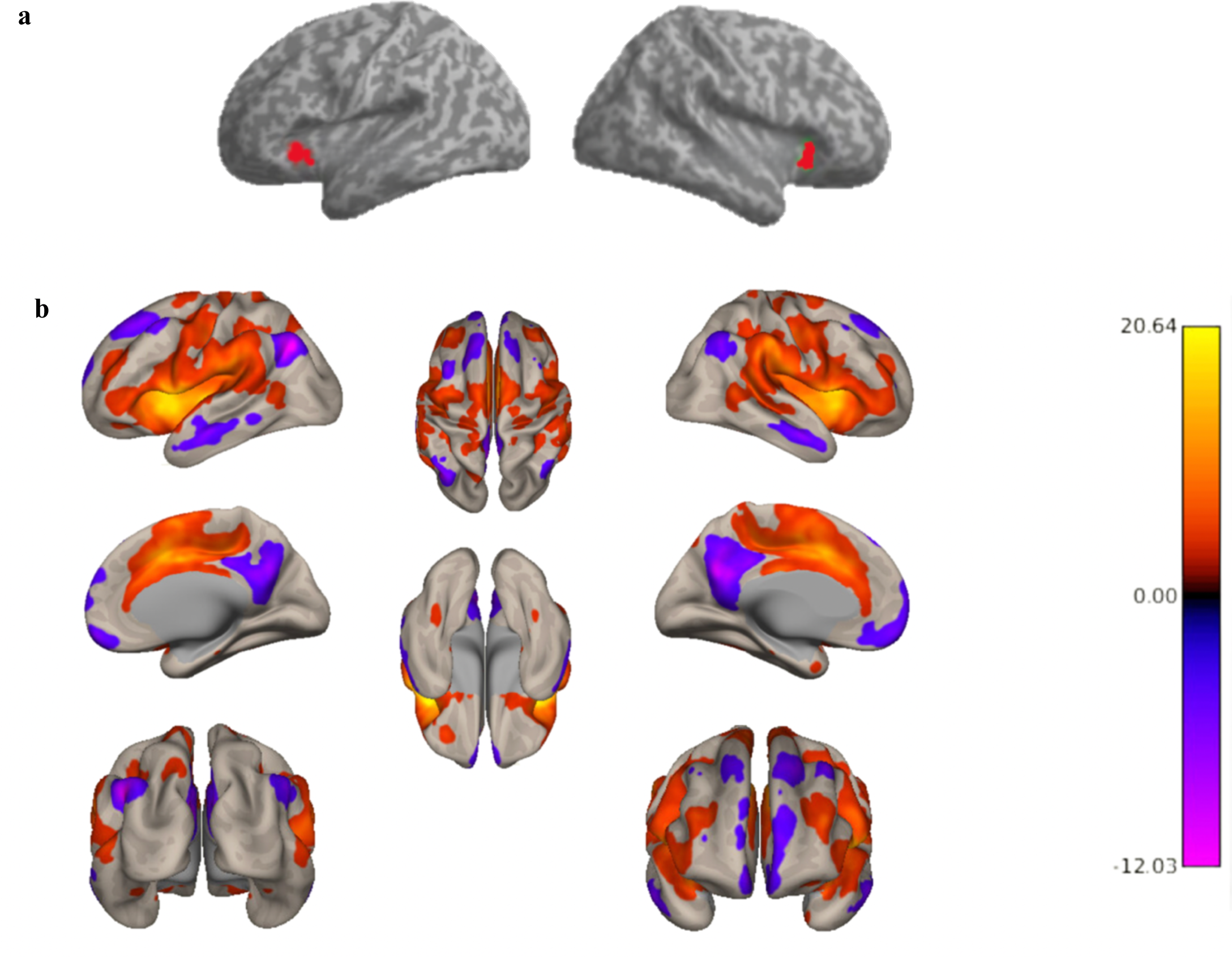
Whole-brain resting-state functional connectivity to the bilateral anterior insula (aI). (a) Illustrates the bilateral anterior insula region of interest (ROI). (b) Summary figure of group-level connectivity from bilateral aI seed ROI showing an expected pattern of insular to whole-brain connectivity: Positive functional connectivity between aI and nodes of the salience network were identified (e.g., anterior cingulate cortex and frontal operculum), in addition to anticorrelations between aI and the default mode network (e.g., medial prefrontal cortex, inferior parietal lobule and precuneus). Connectivity results are overlaid on a MNI template and corrected for multiple comparisons (voxel *p*<0.001; cluster *p*<0.05 FWE-corrected).

## 3 Results

### 3.1 Seed-Based Functional Connectivity Results

#### 3.1.1 PREVENT-AD Sample

To determine whether the anterior insula showed an expected pattern of whole-brain connectivity in our sample, we first conducted a seed-to-voxel analysis for the whole sample without the behavioral covariate of interest. Figure 1b illustrates the functional connectivity pattern from the bilateral anterior insula (aI) to the entire brain for all participants. The expected insular-to-whole-brain connectivity pattern is evident, including positive functional links between the aI and salience network nodes (e.g., anterior cingulate cortex and frontal operculum), as well as inverse or anti-correlations between the aI and nodes of the default mode network (e.g., medial prefrontal cortex, inferior parietal lobule, and precuneus). Next, we performed Generalized Linear Model (GLM) analysis to examine the association between smoking behavior change and the functional connectivity between bilateral aI and the whole brain. The results showed that decreased smoking at follow-up compared to baseline was associated with diminished functional connectivity between bilateral aI and a cluster within the ventromedial prefrontal cortex (vmPFC) that encompasses the following regions: frontal pole, frontal medial cortex, subcallosal cortex, paracingulate gyrus, anterior cingulate cortex, and frontal orbital cortex (Bhanji et al., 2019). Notably, the peak voxel was localized in subcallosal cortex (*F*(2,17) = 19.48; voxel height *p*<0.001 uncorrected; cluster *p*-FWE <0.05; k = 112 voxels; Peak MNI voxel = 0, 30, −8 [x, y, z coordinates]) (Figure 2a). To ensure that our results were not driven by outliers, we performed a post-hoc robust regression analysis to visualize and understand functional connectivity at the level of the individual. We extracted the mean Z-value from the vmPFC cluster for each participant and plotted this against change in smoking across the two timepoints. Findings of the post-hoc analysis showed that our results were not driven by outliers, *r^2^*=0.77, *p*<0.001 (Figure 2b).

**Figure 2.**
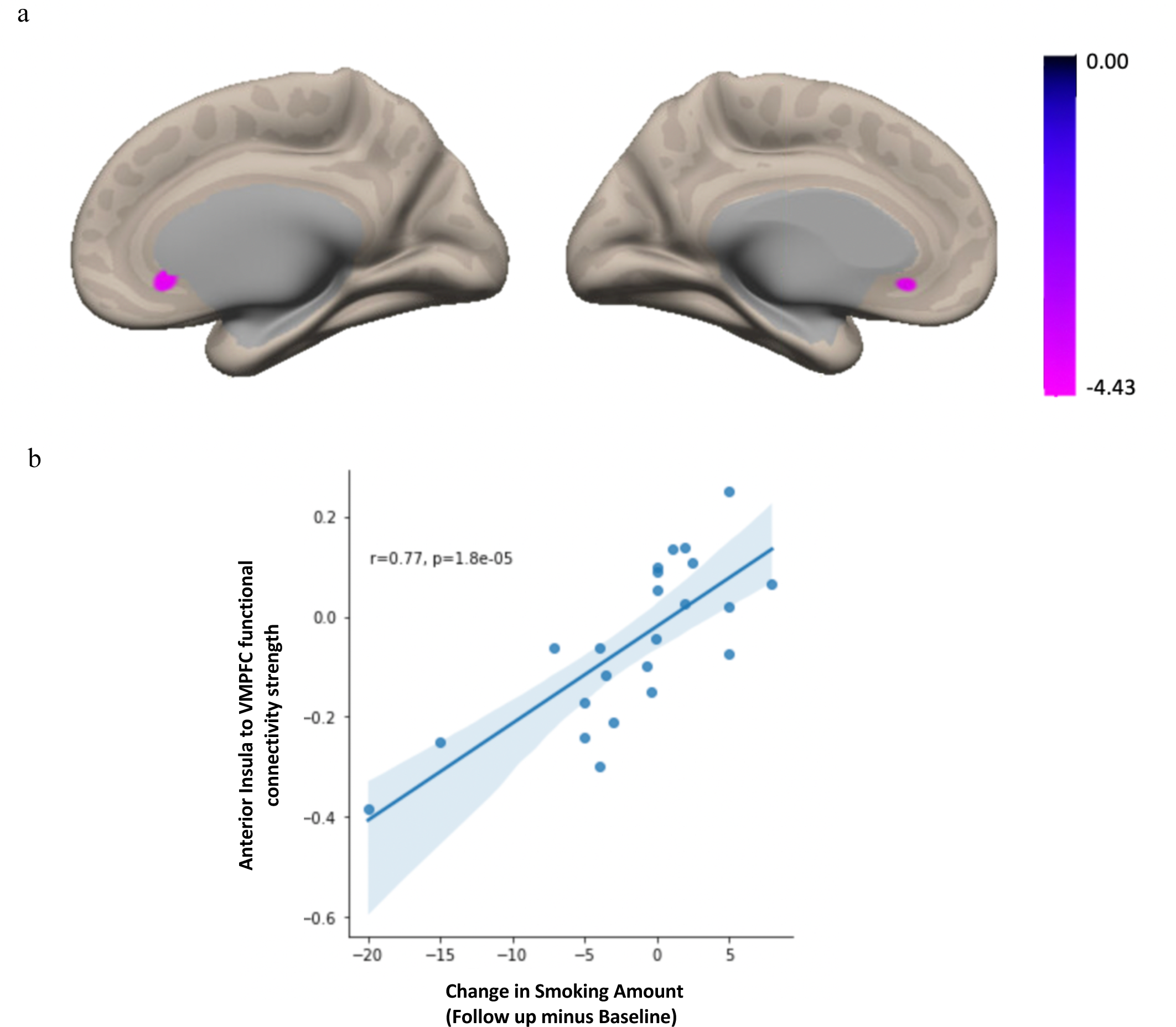
Summary figure of the seed-to-voxel results from the bilateral anterior insula seed in the PREVENT-AD Sample. Decreased smoking at follow-up was associated with diminished functional connectivity between bilateral anterior insula and a cluster within ventromedial prefrontal cortex (vmPFC) (a), with a peak voxel was located within subcallosal cortex. Connectivity results are overlaid on the MNI template brain and corrected for multiple comparisons (voxel *p*<0.001; cluster *p*<0.05 FWE-corrected). Age, sex, baseline smoking amount, *APOE4* carrier status, and mean motion were used as covariates of non-interest. (b) A post-hoc robust regression analysis depicts individual-level connectivity to ensure the results were not driven by outliers.

#### 3.1.2 UK Biobank Preregistered Replication Sample

To replicate and extend the findings of our first study, we examined whether anterior insular connectivity was associated with long-term smoking behavior change in an independent and larger sample from the UK Biobank (Figure 3). The results showed that decreased smoking in follow-up compared to baseline was associated with decreased functional connectivity between bilateral aI and a small cluster within vmPFC with the peak voxel located in the anterior cingulate cortex (ACC), (*F*(2, 17) = 17.48; voxel height p<0.001 uncorrected; cluster p-FWE <0.05; k = 60 voxels; Peak MNI voxel = 0, +18, +24 [x, y, z coordinates]). To again ensure that our results were not driven by outliers, we performed a post-hoc robust regression analysis to visualize and understand functional connectivity at the level of the individual. Excluding outliers over 2.5 SD from mean resulted in the exclusion of 6 participants. We extracted the mean Z-value from the vmPFC cluster for each participant and plotted this against change in smoking across the two timepoints. Findings of the post-hoc analysis showed that our results were not driven by outliers, *r^2^*=0.42, *p*<0.001 (Figure 3b).

**Figure 3.**
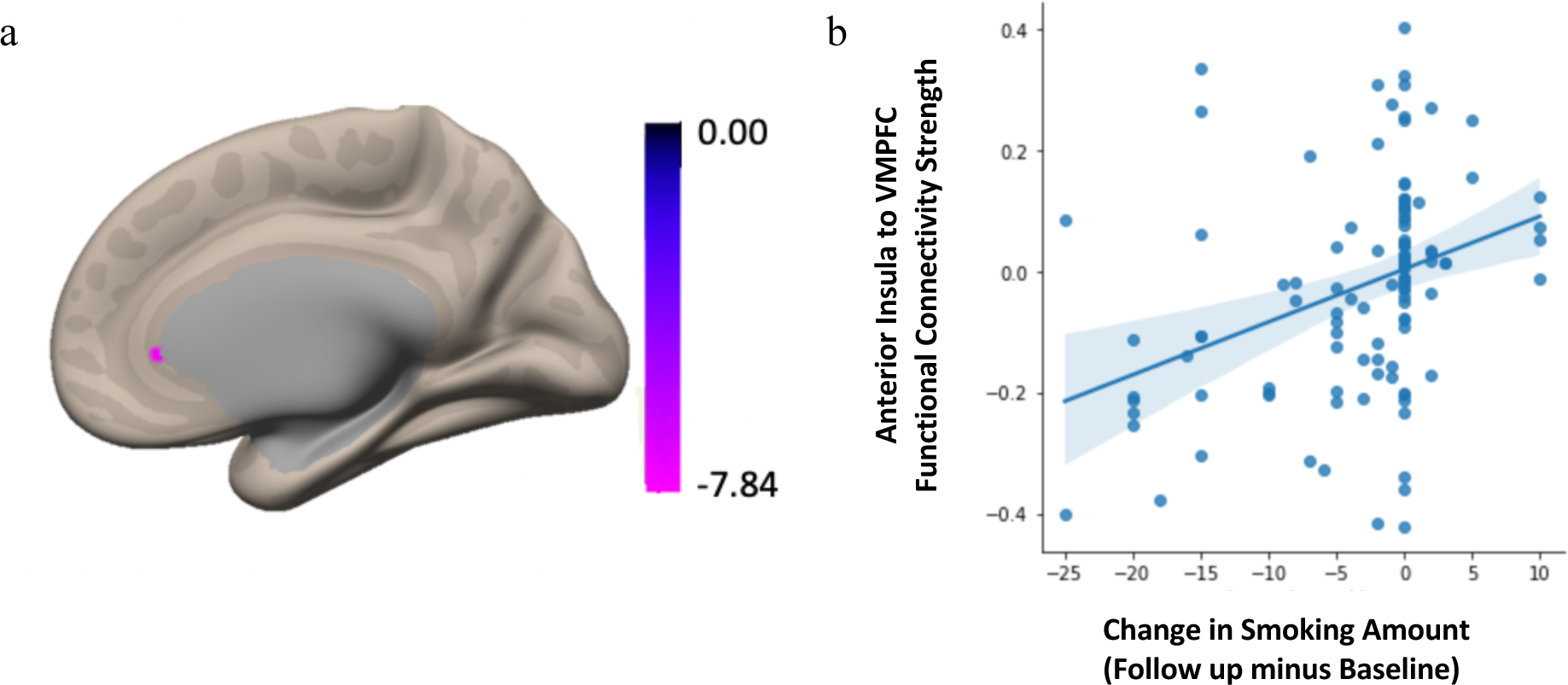
Summary figure of the seed-to-voxel results from the bilateral anterior insula seed in the UK Biobank Replication Sample. Decreased smoking in follow-up compared to baseline was associated with lower functional connectivity between bilateral anterior insula and a cluster within the ventromedial prefrontal cortex (vmPFC) (a), with a peak voxel located within the anterior cingulate cortex. These connectivity results are overlaid on the MNI template brain and corrected for multiple comparisons (voxel *p*<0.001; cluster *p*<0.05 FWE-corrected). Age, sex, baseline smoking amount, *APOE4* carrier status and mean head motion were included as covariates of no interest. (b) A post-hoc analysis allowed visualization of individual-level aI-VMPFC functional connectivity strength to ensure that the main results were not driven by outliers.

## 4 Discussion

The primary goal of the current study is to characterize the neural substrates associated with long-term smoking reduction in typical and at-risk aging. We conducted a hypothesis-driven rs-fcMRI seed-to-voxel analysis in a primary sample of current smokers who are cognitively normal older adults at an increased risk for AD from the PREVENT-AD cohort. We identified that diminished baseline functional connectivity between anterior insula (aI) and ventromedial prefrontal cortex (vmPFC) involving frontal pole, frontal medial cortex, subcallosal cortex, paracingulate gyrus, anterior cingulate cortex (ACC), and frontal orbital cortex, with a peak voxel in the subcallosal cortex, was associated with decreased smoking in long-term follow-up after 2.7 years. The robustness of our results was confirmed through a pre-registered replication in a larger sample of cognitively normal older adults who were current smokers from the UK Biobank longitudinal cohort. Both studies applied identical preprocessing and analytic approaches. In the replication study, we found that smoking maintenance behavior was again associated with differential frontoinsular connectivity: Decreased smoking at long-term follow up after 5.2 years was associated with decreased aI-vmPFC functional connectivity, with a peak voxel in ACC. This supports the validity and generalizability of our results and underscores the importance of aI-vmPFC functional connectivity in long-term smoking reduction maintenance in typical and at-risk aging.

Our central findings can be understood in light of prior research implicating coordinated activity between anterior insula and ACC in goal-directed behavior and decision-making (Craig, 2002; Fedota et al., 2018; Janes et al., 2010, 2015; Menon & Uddin, 2010; Uddin, 2015). In line with these roles, the anterior insula is critical for awareness of bodily urges, while the ACC is important in initiating behaviors and resolving conflict (Craig., 2002). Prior evolutionary research provides context for the coordinated activity between these regions; ACC evolved first as a motor-control region aligned with the sensory integration of olfactory-guided group behavior in mammals, while the insula evolved later for cortical processing of homeostatic sensory activity in the individual animal (Heimer & Van Hoesen, 2006). Their conjoint activation is therefore thought to constitute a system of self-awareness: conscious interoception arising from the anterior insula is re-represented in the anterior cingulate cortex, forming the basis for selecting and preparing responses to inner or outer events (Allman et al., 2011; Janes et al., 2015; Medford & Critchley, 2010). The integration of interoceptive representations is thought to progress sequentially from the posterior insula to the anterior insula (Craig et al., 2000; Olausson et al., 2002) The primary interoceptive stimuli, encompassing numerous individually mapped and distinct feelings from the body, are established in the posterior insular cortex (Craig, 2002; Craig et al., 2000). This information is then transmitted to the anterior insula, where re-represented interoceptive signals are integrated with emotional, cognitive, and motivational signals from other cortical and subcortical regions, such as the amygdala, the anterior cingulate cortex, the dorsolateral prefrontal cortex and the ventral striatum (Brooks et al., 2002; Craig, 2002; Craig et al., 2000).

The role of insula and vmPFC in reward-related decision making and thereby, smoking behaviour change is well established (Bechara et al., 2002; Bechara & Damasio, 2002; Mitchell, 2004; Spinella, 2002). VMPFC is a node in the reward and default mode networks that is relatively spared in aging and AD, integrates value, social and self-referential processing, and is linked to successful behaviour change (Alexander et al., 2023; Damasio et al., 2000; A. H. Gutchess et al., 2007; A. Gutchess & Samanez-Larkin, 2019; Kang et al., 2018; Naqvi et al., 2007; Salat et al., 2005; Winecoff et al., 2013). For example, a study that applied graph theory and modularity analyses showed that higher vmPFC flexibility (i.e., the degree to which the vmPFC changes its community assignment over time) is positively correlated with smoking behavior change a month later (Cooper et al., 2018). Our current findings support the existing body of work suggesting aI-vmPFC regions work in concert to maintain smoking change behaviour.

Prior lesional and fMRI studies have lead to seemingly contradictory findings about the role of insula in smoking behavior (Droutman et al., 2015). On one hand, numerous functional neuroimaging studies have demonstrated reduced activity in the insular cortex among individuals with substance dependence during decision-making tasks (Nestor et al., 2010; Stewart, Connolly, et al., 2014; Stewart, May, et al., 2014). Lower insular cortex activation was shown to be predictive of relapse after a period of abstinence (Droutman et al., 2015). This work suggests diminished cognitive control, as indexed by lower activation of insular cortex, in individuals at risk of relapse. In contrast, evidence from lesion studies presents a paradoxical scenario wherein nicotine-addicted individuals have been able to quit smoking abruptly and effortlessly following damage to the insular cortex (Naqvi et al., 2014). Lesional studies highlight the role of diminished insular recruitment leading to reducing cigarette cravings. Towards a systems-level understanding, some fMRI studies suggest that decreased insular connectivity with DMN regions, including vmPFC ameliorates nicotine withdrawal symptoms and is associated with decreased smoking (Janes et al., 2020; Sutherland et al., 2013). Our findings align with the latter perspective and evidence from lesional work. Teasing apart the potential contribution of component cognitive processes such as emotion regulation, temporal discounting, cognitive control from the role of cravings and urges will be an important future direction of this work.

The current findings are in line with the framework of Incentive Sensitization Theory (IST). IST posits that drug addiction is characterized by a hyper-reactive mesolimbic dopaminergic system that responds to rewarding cues, resulting in heightened “wanting” to consume more drugs (Berridge & Robinson, 2016; Olney et al., 2018). This phenomenon, known as incentive salience, refers to the attention-grabbing and motivational features of rewards and their associated cues (Bindra, 1978). Reward cues have the ability to trigger bursts of reward-seeking motivation and can become a “motivational magnet” over time (Olney et al., 2018; Peciña & Berridge, 2013; Robinson et al., 2015). The incentive salience circuitry is mediated by larger brain systems involving mesocorticolimbic dopamine, the central amygdala (CeA), lateral hypothalamus (LH), orbitofrontal cortex (OFC), ventromedial prefrontal cortex and the insular cortex (Castro et al., 2015; DiFeliceantonio & Berridge, 2016; Naqvi & Bechara, 2010; Olney et al., 2018; Warlow et al., 2017). In line with IST, disruption of frontoinsular circuitry would be expected to reduce incentive salience or ‘wanting’ with subsequent reduction in smoking behavior.

Indeed, Naqvi et al., (2010) proposed a model for insula involvement in drug addiction-related decision making: The insula plays a key role in weighing the positive and negative consequences of addictive substances. According to the model, the insula encodes the positive hedonic effects and the physical sensations experienced during drug use, which are recalled during drug cravings through coordinated activity with the amygdala, orbitofrontal cortex (OFC), vmPFC and dorsolateral prefrontal cortex (DLPFC). This can manifest as an urge to use the drug. Simultaneously, the insula represents the negative consequences of drug use in terms of its impact on bodily integrity, survival and homeostasis. The anterior cingulate cortex is responsible for detecting conflicts between the goal of drug use and other goals and shifting attention accordingly. Therefore, the decision to abstain depends upon the ability to suppress the representation of the positive hedonic effects of drug use and to enhance the representation of the negative consequences of drug use. This suggests that successful and sustained smoking cessation occurs when the representation of the negative consequences of drug taking outweighs the representation of the positive hedonic effects of drug taking. Insular lesions might support successful behaviour change as the hedonic impact would be significantly diminished.

Evaluation and support for the model, however, has largely been limited to lesion studies and acute smoking abstinence behavior, over only a few days to months.

Smoking might exert different effects on insular connectivity depending on the AD risk level – be it high or low. Past MRI investigations have showcased loss of insular gray matter (Guo et al., 2012) and disrupted insular networks in the early stages of AD (Xie et al., 2012).

These studies have underscored the significance of the insula, which emerges as a potentially vulnerable region and a pivotal hub in AD patients. A recent study explored the potential effects of smoking on insula functional connectivity among patients with mild cognitive impairment (MCI) and found that anterior insular connectivity decreased among smokers with MCI (Zhang et al., 2023).

Our study provides support for expanding the investigation of neurocognitive processes underlying sustained behavior change to include smoking reduction, in addition to cessation.

Typically, smoking outcomes are dichotomized in studies as quitters versus non-quitters. However, any reduction in smoking behavior, even if not leading to complete cessation, can still have positive effects on cognitive functioning and lower the risk of dementia (Begh et al., 2015). Moreover, individuals who successfully reduce their smoking are more likely to attempt and achieve complete cessation (Balfour, 2009). Thus, examining the neural substrates of smoking reduction provides valuable mechanistic insights with translational potential in individuals at risk for AD.

The most compelling evidence for the critical role of insula in maintaining drug-related urges comes from human lesion studies: Acute insular injury abruptly disrupts cigarette cravings and produces acute smoking cessation (Naqvi et al., 2007). Future studies combining functional neuroimaging and lesional methods will help strengthen the causal cascade between insular injury and diminished aI-vmPFC functional connectivity. Additionally, exploring effective connectivity would offer valuable insights into the directionality of the insular connectivity relationship with smoking cessation behavior, making this an intriguing avenue for further investigation.

It is important to acknowledge that a key limitation of the current study is the correlational nature of functional connectivity analyses which do not allow for determination of causality. Additionally, our replication cohort (UK-Biobank) significantly differs in its composition when compared to our primary discovery cohort, PREVENT-AD. Notably, PREVENT-AD consists mainly of French-Canadian participants, a demographic distinction that does not hold true for the UK-Biobank dataset. Furthermore, the UK-Biobank is not specifically designed to study individuals at risk for AD. Similarly, the PREVENT-AD dataset was not explicitly designed to explore the effects of smoking, and as such detailed smoking data, beyond that of self-reported smoking status, is lacking. Taken together, further replication is needed in diverse cohorts to validate our findings.

Despite these limitations, our study has important strengths, including the use of a longitudinal sample of older adults at-risk for AD and a rigorous and hypothesis-driven approach in a pre-registered larger replication and extension study. The translational potential of our findings include the identification of the aI-vmPFC circuit as a potential therapeutic target for smoking cessation strategies. Neurostimulation approaches to behavior change are in their infancy with some early evidence in physical activity engagement in aging (Lo et al., 2021) as well as in smoking cessation (Dinur-Klein et al., 2014). Furthermore, our study highlights the importance of a brain-based mechanistic approach to understanding behavior change and resilience in typical and at-risk aging.

## Data and Code Availability

Publicly available software used for all the analyses is CONN toolbox (https://web.conn-toolbox.org/). The individual-level UK Biobank data can be obtained from https://www.ukbiobank.ac.uk/. Requests to access the PREVENT-AD dataset used in this study can be made to the StoP-Alzheimer team: https://prevent-alzheimer.net/?page_id=1760&lang=en.

## Author Contributions

N.T: Conceptualization, Methodology, Formal analysis, Investigation, Data curation, Writing— original draft, Writing—review & editing, Visualization. M.I: Investigation, Writing—review & editing. A.N.C., and C.W: Data curation, Writing—review & editing. G.B: Methodology, Data curation. J.T.M: Methodology, Data curation, Resources, Writing—review & editing. S.V: Methodology, Resources, Writing—review & editing. R.N.S: Methodology, Resources, Data curation. M.R.G: Conceptualization, Methodology, Resources, Writing—review & editing, Funding Acquisition, and Supervision.

## Funding

This Project has been made possible by a National Sciences and Engineering Research Council of Canada (NSERC) Discovery Grant (DGECR-2022-00299), an NSERC Early Career Researcher Supplement (RGPIN-2022-04496), a Québec Brain Imaging Network Grant, a Fonds de Recherche Santé Quebec (FRSQ) Salary Award, the Canada Brain Research Fund (CBRF) an innovative arrangement between the Government of Canada (through Health Canada) and Brain Canada Foundation and the Alzheimer’s Society of Canada New Investigator Grant to MRG. This research was undertaken thanks also to funding from the Canada First Research Excellence Fund, awarded to the Healthy Brains Healthy Lives initiative at McGill University to NT.

## Declaration of Competing Interest

The authors declare no competing interests.

## Ethics Statement

This study utilized data from the UK-Biobank study and the PREVENT-AD study. The UK-Biobank has obtained ethics approval from the North West Multi-Centre Research Ethics Committee (MREC, approval number: 11/NW/0382), and obtained written informed consent from all participants prior to the study. All protocols and study procedures in the PREVENT-AD study were approved by the McGill Institutional Review Board and/or Douglas Mental Health University Institute Research Ethics Board in accordance with the Declaration of Helsinki.

## Acknowledgements

We are extremely grateful to the research participants who dedicated their time to participating in this study. We would also like to thank members of the PREVENT-AD and UK Biobank Research Groups. Finally, we would like to thank lab members who participated in data collection for the PREVENT-AD study, including Maggie Nguyen, Sarah Elbaz, Andréanne Powers, Linda Li, Sarah Elbaz, Helen Pallett-Wiesel, Nicolas Lavoie and Shania Fock Ka Bao.

## Notes

https://www.ukbiobank.ac.uk/

https://prevent-alzheimer.net/?page_id=1760&lang=en

### Competing Interest Statement

The authors have declared no competing interest.

### Summary of Updates

Author list order updated

